# The faecal microbiome of the wild European badger *Meles meles*; a comparison against other wild omnivorous mammals from across the globe

**DOI:** 10.1101/2022.03.17.484750

**Authors:** James Scott-Baumann, Jessica C A Friedersdorff, Bernardo Villarreal-Ramos, Jonathan King, Beverley Hopkins, Richard Pizzey, David Rooke, Glyn Hewinson, Luis A. J. Mur

## Abstract

Here we investigate the faecal microbiome of wild European badgers *Meles meles* using samples collected at post-mortem as part of the All Wales Badger Found Dead study based on sequencing the V3-V4 region of the 16S rRNA gene. This is the first published characterisation of the badger microbiome. We initially undertook a sex-matched age comparison between the adult and cub microbiomes. Analysis used the QIIME 2 pipeline utilising DADA2 and the Silva database for taxonomy assignment. Fusobacteria appeared to be more abundant in the microbiomes of the cubs than the adults although no significant difference was seen in alpha or beta diversity between the adult and cub badger microbiomes. Comparisons were also made against other wild, omnivorous, mammals’ faecal microbiomes using publicly available data. Significant differences were seen in both alpha and beta diversity between the microbiomes from different species. As a wildlife species of interest to the disease bovine tuberculosis, knowledge of the faecal microbiome could assist in identification of infected badgers. Our work here suggests that if comparisons were made between the faeces of bTB infected and non-infected badgers, its possible age may not have a significant impact on the microbiome.

## Introduction

Bovine tuberculosis (bTB) poses a huge economic cost to UK cattle farming (Defra, 2020). Badgers are known to carry the causative organism, *Mycobacterium bovis*, and can pass it to farmed cattle (Godfray *et al.*, 2018). Current tests available for diagnosis of bTB in badgers are limited by sensitivity and practicality (requiring trapping live animals) (Thomas and Chambers, 2021). In humans, changes in the gut microbiome have been shown to occur with tuberculosis infection (Luo *et al.*, 2017). Although tuberculosis is a pulmonary disease, and changes in the gut microbiome secondary to changes in the lung microbiome it in is now referred to as the gut-lung axis (Enaud *et al.*, 2020). These changes in the gut microbiome may hold potential diagnostic purposes (Hu *et al.*, 2019) that could then be used on faecal samples easily collected from around setts, avoiding trapping of live animals. However, before assessing the badger’s faecal microbiome changes associated with bTB infection, there must be an understanding of the healthy microbiome and the associated sources of variation. Age has been shown to the one of the most significant factors affecting the microbiome of a species, with maturation of the microbiome over time from birth to adulthood (Conlon and Bird, 2014). In this study, a comparison is first made between the faecal microbiomes of adult and cub (’1 year old) badgers collected during post-mortem as part of the ‘All Wales Badger Found Dead’ project. Secondly, a comparison is made between these badger microbiomes and 24 other faecal microbiomes from different wild, omnivorous mammal species collected from the environment from across the globe (Youngblut et al. 2019).

## Results and Discussion

### Intra-species age comparison of badger microbiomes

Faecal samples were taken from the rectum of wild badgers found in Wales and known to be *M. bovis* negative through culture. Following DNA extraction using a FastDNA SPIN kit for soil (MP Biomedical, Santa Ana, USA) the Illumina MiSeq platform was used to amplify the V3-V4 region of the 16S rRNA (https://support.illumina.com/documents/documentation/chemistry_documentation/16s/16s-metagenomic-library-prep-guide-15044223-b.pdf). All downstream analysis of the raw read files was done using the QIIME 2 pipeline (QIIME2 v2021.4, Bolyen et al., 2019). Primers were trimmed from reads, and forward and reverse reads were trimmed when PHRED score dipped below 20. Sampling depth was cut-off at 3816 reads in order to keep the sample with the lowest number of reads (fat dormouse). Rarefaction curves suggested minimal loss of diversity at this cut-off (Figure S1). Taxonomy was assigned to OTUs using the Silva database (Quast et al., 2013; Bokulich et al., 2018; Robeson et al., 2020).

More than 99.9% of the operational taxonomic units (OTU) were assigned to known phyla using the SILVA database for both the adults and the cubs. The most predominant bacterial phyla present in the badgers’ faecal microbiomes were Proteobacteria and Firmicutes, together accounting for 75% or more of the percentage abundance across all samples (Figure 1). This contrasted with other human and animal studies which have often shown a typical predominance of Firmicutes and Bacteroidetes (Eckburg et al., 2005; Proudman et al., 2014; Brice et al., 2019; Shanmuganandam et al., 2020). This greater predominance of Proteobacteria and Firmicutes, with a lower proportion of Bacteroidetes, is more similar to the basic patterns that have been found in wild and captive birds (Grond *et al.*, 2018) and some marine mammals (Nelson *et al.*, 2015). Fusobacteria appeared to be more abundant in the microbiomes of the cubs (mean ~7.5%) than the adults (mean ~0.1%). The “kit-ome” blank sample showed a wider diversity of phyla suggesting that this was not a source of contamination bias in the badger faecal microbiomes.

**Figure 1.**
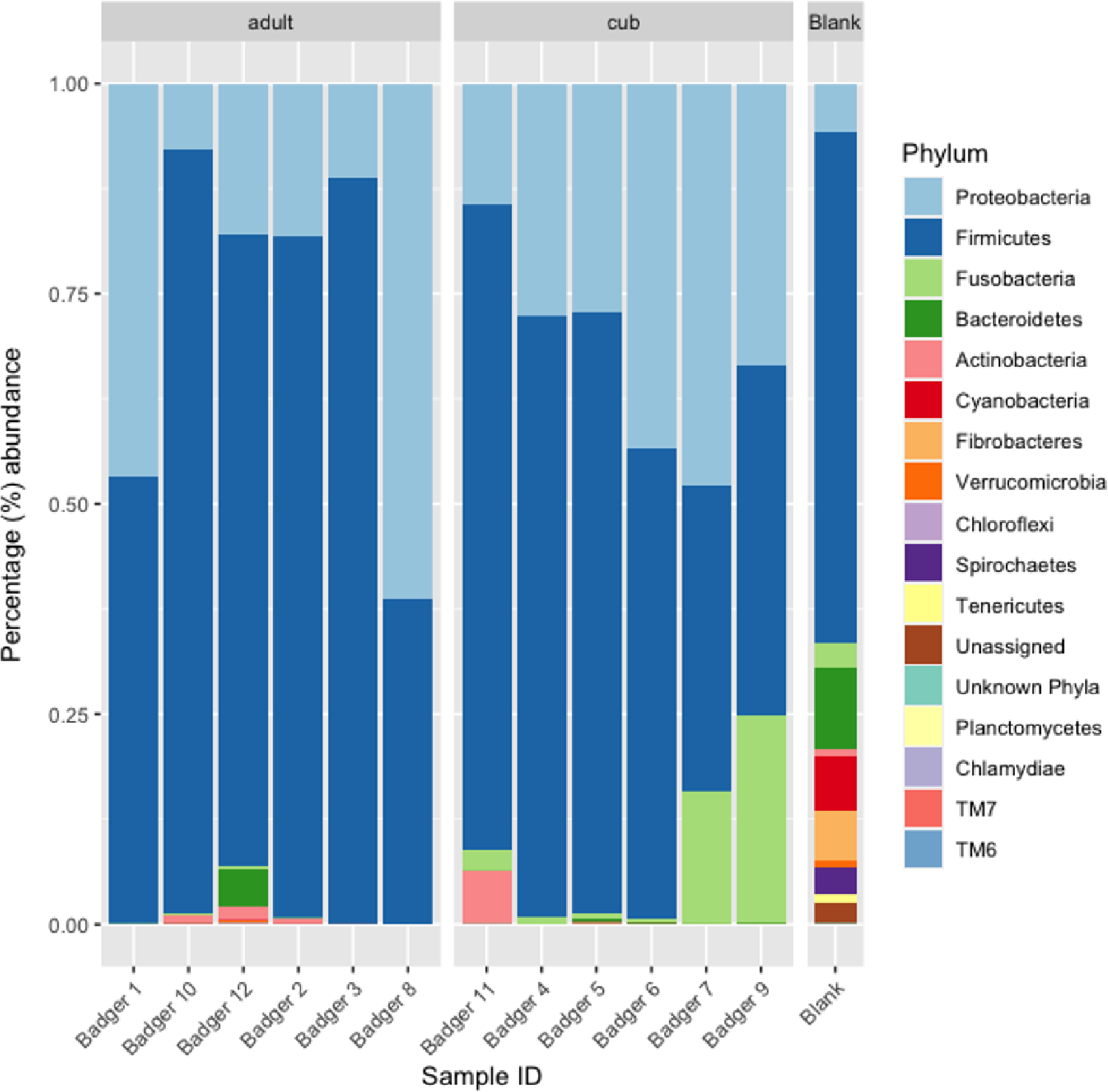
Percentage abundances of different phyla of bacteria present in the faecal microbiome of adult and cub badgers

At the genus level the proportion of OTUs assigned for adults and cubs were 68.9% and 54.2% respectively. At the species level only 7.7% and 6.5% of OTUs were linked to uncultured species or those identified from previous metagenomes, for adults and cubs respectively. Of the OTUs successfully identified using SILVA taxonomy, *Romboutsia hominis*, and genera *Shigella*, *Clostridium sensu stricto 1*, *Paeniclostridium* and *Terrisporobacter* were common to all badger samples. There were no species or genera found to be unique to all cub or to all adult samples.

Alpha diversity comparisons (Shannon and Simpson’s index) showed no significant differences (*p*=0.42 each) between the two age groups using Kruskal-wallis pairwise (Figure 2a). Beta Diversity comparisons (Bray-Curtis, Jaccard, unweighted Unifrac, weighted Unifrac) showed no significant differences (*p*=0.14, 0.71, 0.19, 0.13 respectively) between the two age groups using PERMANOVA (Figure 2b). This indicated that the adult and cub badgers’ faecal microbiomes were similarly diverse in our sample population.

**Figure 2.**
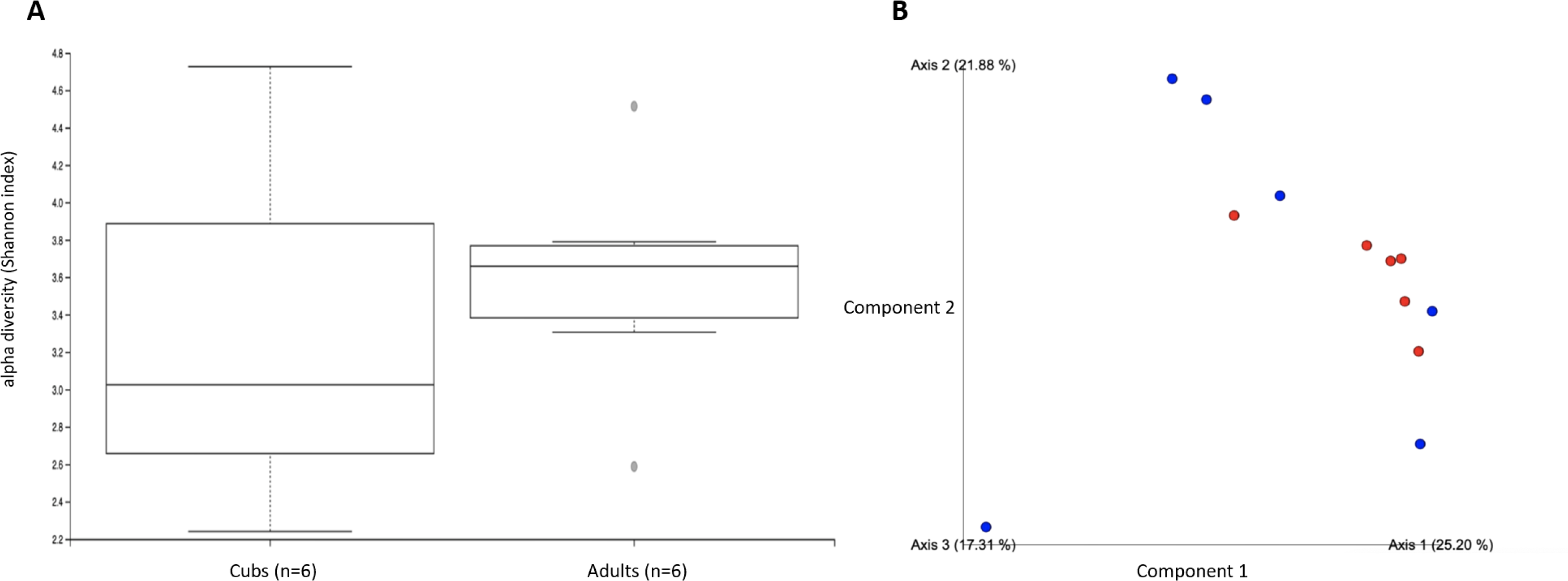
Diversity estimates of the badger microbiome based on (A) alpha diversity (Shannon index) and (B) beta diversity (Bray-Curtis dissimilarity) comparing cubs (red) and adults (blue).

Age has previously been shown in both human and animal models to be a significant factor in the faecal microbiome, with its development and maturation over time (Conlon & Bird, 2014; Lees et al., 2014; Deusch et al., 2015). Given that the only cubs recruited into this study were old enough to be free roaming and leave the sett, they may have already developed a ‘mature’ microbiome, when compared to those cubs that are young enough to reside in the setts, which would not be captured in this study by the nature of the sampling methods. Knowing whether a faecal microbiome changes with age is key when going on to look for marker species present in faeces as an indicator for bTB infection.

### Inter-species comparison of wild omnivore microbiomes

To provide a comparison of faecal microbiomes from other wild mammals, samples were downloaded that were generated from the Youngblut et al. (2019) study (European Nucleotide Archive, study accession number PRJEB29403). All those used were faecal microbiomes from omnivorous, wild mammals (n=24) and were generated using primers for just the V4 region of the 16S rRNA. The percentage abundances of all bacterial phyla identified across all samples shows that Firmicutes, Proteobacteria, Fusobacteria and Bacteroidetes are the most prevalent (Figure S2). Alpha diversity comparisons showed that badger faecal microbiomes were more diverse than other members of the family Mustelidae (beech marten, pine marten), but were less diverse than other mammals such as the European ground squirrel, which had the highest alpha diversity. There were significant differences between faecal microbiomes at the order (*p*=0.005), family (*p*=0.005) and genus (*p*=0.04) levels of mammal host groupings with the Shannon diversity metric (Figure 3). Beta Diversity comparisons (Bray-Curtis, Jaccard, unweighted Unifrac, weighted Unifrac) all showed significant differences between the different hosts at the level order (p<0.001), family (p<0.001) and genus (p<0.001) using PERMANOVA (Figure 4). Differences the percentage abundance between bacterial phyla identified in the microbiomes are displayed using hierarchical clustering (Figure 5).

**Figure 3:**
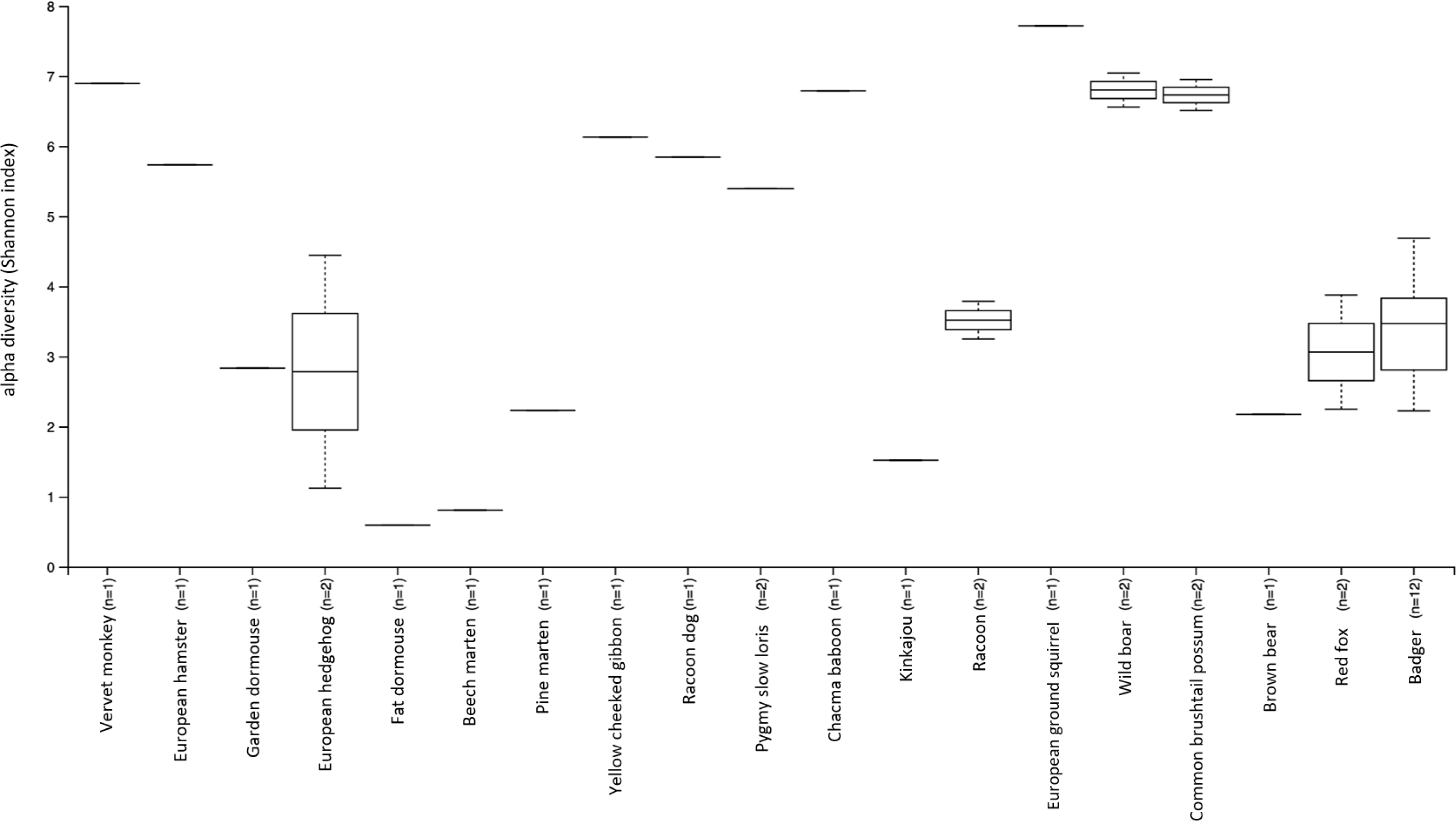
Assembling the omnivorous mammalian faecal microbiome collection: Alpha diversityusing the Shannon index

**Figure 4:**
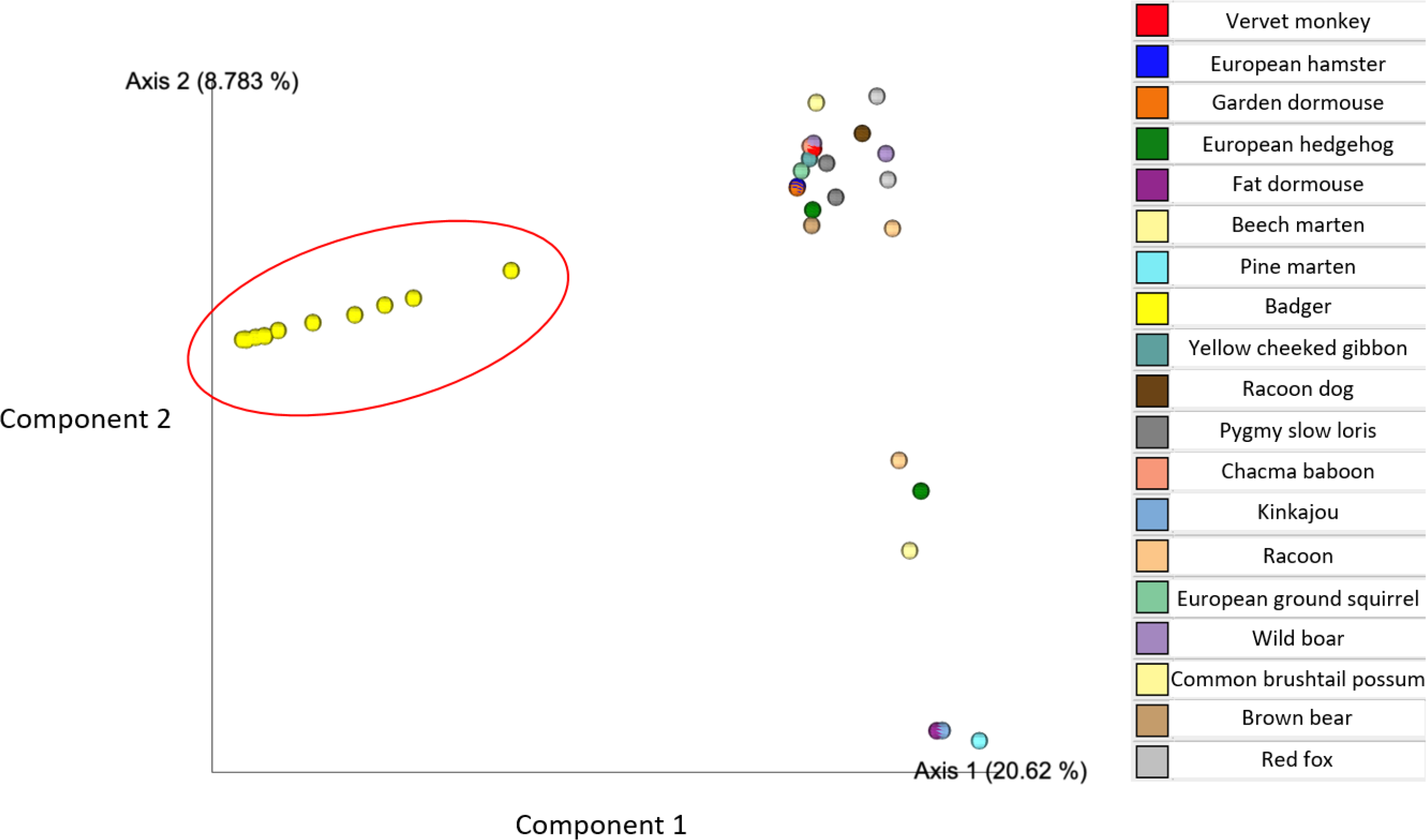
Assembling the mammalian faecal microbiome collection: Principal coordinates analysis showing beta diversity, based on the Bray-Curtis dissimilarities. European badgers *Meles meles* (yellow, n=12) separate clearly as a group

**Figure 5:**
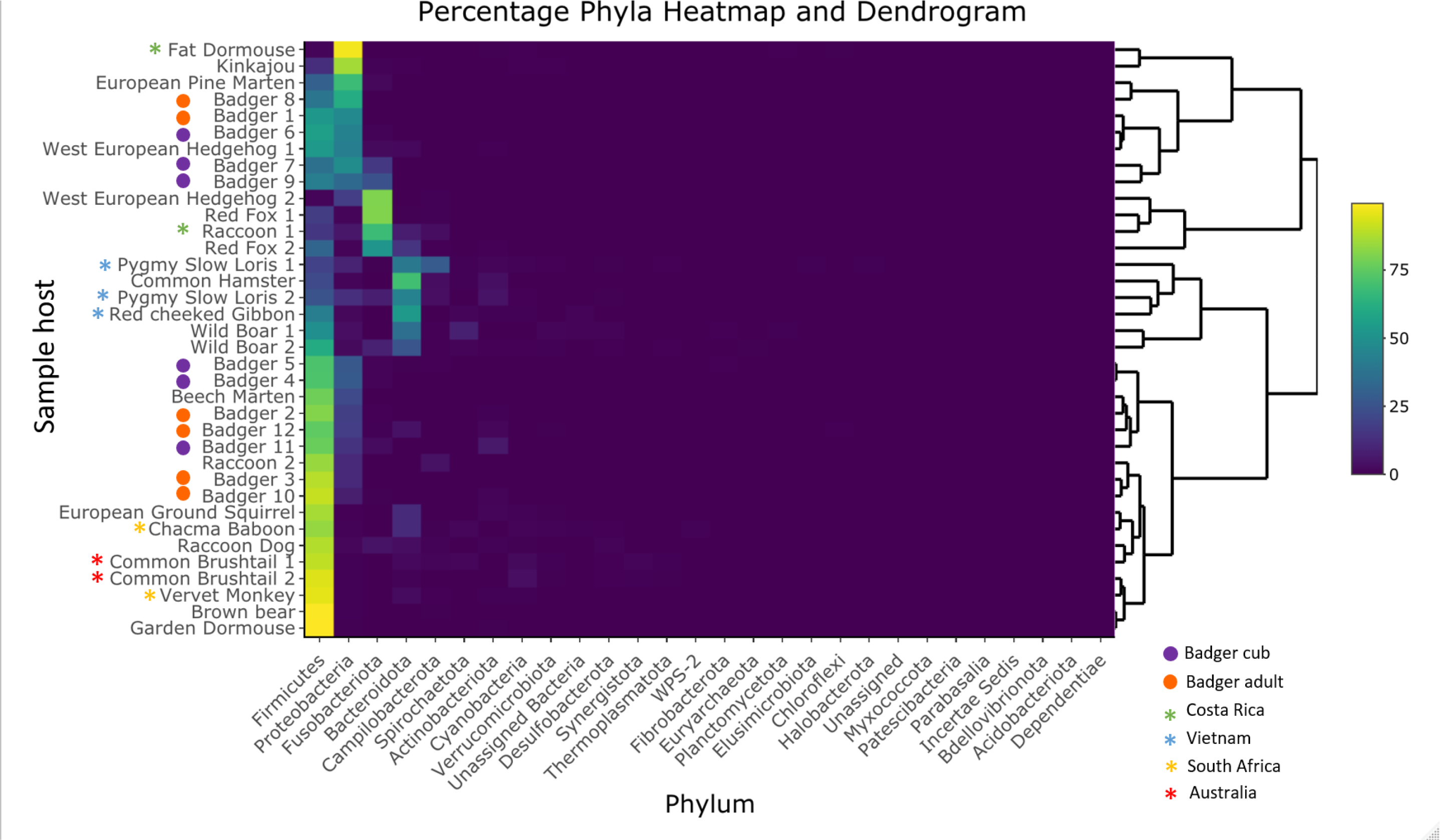
Dendrogram created from percentage abundance of bacterial phyla present in their faecal microbiomes. Colours in the heatmap correspond to the legend on the right, with yellow 100% and purple 0% percentage abundance. Asterisks denote country sampled from.

Several studies have shown that host phylogeny and diet have significant impact on the faecal microbiome (Ley *et al.*, 2008; Colston and Jackson, 2016), and captivity has also been shown to affect the microbiome markedly (Clayton *et al.*, 2016; Gibson *et al.*, 2019; Schmidt *et al.*, 2019; Ning *et al.*, 2020). From the Youngblut et al. (2019) study only hosts that were omnivorous, wild and mammalian were selected, thereby allowing closer comparisons with the badgers. These 24 samples showed a predominance of Firmicutes followed by Bacteroidetes, which is in line with several other studies (Eckburg *et al.*, 2005; Proudman *et al.*, 2014; Brice *et al.*, 2019; Shanmuganandam *et al.*, 2020), compared to badgers. This could be influenced by the badger samples being collected from the rectum of deceased animals which may have spent up to 4 days in cold storage. This time in cold storage was also additional to the time carcasses spent in the environment before they were seen and collected. These samples were in contrast to the other wild mammal samples that were collected as faecal samples from the field. Pechal *et al.* (2018) found that post-mortem microbiome changes in humans were significantly different only after 48 hours after death. Considering the impact of refrigeration, Choo *et al.*, (2015) found that refrigeration of faecal samples at 4°C for 72 h before 16 rRNA analysis did not differ significantly from those stored immediately at −80°C.

Considering the differences between the faecal microbiomes (Figure 5), the influence of diet is likely to be a major determinant. One particular group of host species, which includes the two red foxes, one West European hedgehog and one racoon, had higher levels of Fusobacteria, and formed a clade (see Fusobacteroita, Figure 5). Several studies, both human and animal, have shown a proportional decrease in the ratio of Fusobacteria to Firmicutes associated with a higher fibre diet (Middelbos *et al.*, 2010; Hooda *et al.*, 2012; Panasevich *et al.*, 2015). Cats fed diets with a lower protein content have also been shown to have significantly lower levels of Fusobacteria in their faecal microbiome (Hooda *et al.*, 2013). This may be associated with the fact that species like *Fusobacterium varium* can catabolise amino acids as well as carbohydrates (Potrykus *et al.*, 2007). Red foxes, depending on what food is available with regards to the season, local habitat and proximity to humans, can be almost exclusively carnivorous (Sidorovich *et al.*, 2006). West European hedgehogs can similarly be almost entirely insectivorous in some situations (Gimmel *et al.*, 2021). Racoons are highly opportunistic and their proximity to humans and access to anthropogenic food can affect their diet, so much so that it has resulting effects on their health (Hungerford *et al.*, 1999; Schulte-Hostedde *et al.*, 2018). One could hypothesise therefore that the animals in this group might have adopted a more carnivorous diet, which led to greater levels of Fusobacteria in their microbiome.

Another group, including both the wild boar, both pygmy slow loris, the red-cheeked gibbon and the common hamster, had much higher levels of Bacteroidetes than the rest of the samples. In both human and animal models Bacteroidetes can be altered with dietary changes; being positively associated with fat but negatively associated with fibre (Wu *et al.*, 2011; Heinritz *et al.*, 2016). Wan et al. (2019) also found an increased abundance of Bacteroidetes with a high fat diet, with a simultaneous decrease in Firmicutes. However another study found increased proportions of faecal *Bacteroides* associated with a high carbohydrate/high glycaemic index (rapidly digested) diet, rather than high fat diets (Fava *et al.*, 2013). Diet is clearly is significant factor for faecal microbiomes, and the diet of badgers has been shown to vary greatly, even within individuals in the same sett who therefore have access to the same dietary resources (Robertson *et al.*, 2014). This could therefore explain the variability seen in the badgers’ microbiomes here.

Considering alternative influencing factors, the microbiomes could be influenced by geographical origins of sample. Most faecal samples came from Europe, a few coming from South Africa, Australia, Costa Rica and Vietnam. Figure 5 suggests that origins could influence the distribution of percentage abundance of Phyla present in animals’ faecal microbiomes. For example, all samples from Vietnam (the red-cheeked gibbon and the two pygmy slow loris samples) were closely located. Similarly, the microbiomes of the common brushtail possums (both from Australia) and the chacma baboon and vervet monkey (both from South Africa) appeared close together. The two raccoon samples, which are the only replicate samples (from the same species) that are not closely associated on the dendrogram, are from separate continents; Costa Rica (Racoon 1) and Austria (Raccoon 2).

Care must be taken when drawing conclusions about the individual microbiomes here and especially interpreting them as being representative of that species, given that for most host species from the Youngblut et al. (2019) study they were presented by a single faecal sample from a single individual. The importance of this is indicated by the variation seen in our multiple assessments of badger microbiomes (n=12). Indeed, the microbiome of some species can vary much more widely between individuals, when compared to other closely related species; such as hares and rabbits (Shanmuganandam *et al.*, 2020).

This study is the first of its kind published on the faecal microbiome of wild European badgers. Despite the possible limitations of using post-mortem samples, it provides an initial understanding of the faecal microbiome for this population at post-mortem. Multiple studies have used post-mortem samples from badgers for monitoring of bTB (Abernethy *et al.*, 2011; Goodchild *et al.*, 2012; Sandoval Barron *et al.*, 2018; Schroeder *et al.*, 2020) and this work provides evidence that such samples could also be used for microbiome analysis, for instance in the comparison of *M. bovis* infected and non-infected badgers.

## Abbreviations

bTB: bovine tuberculosis

## Acknowledgements

We would like to acknowledge Dr. Matt Hegarty and Dr. Charly Morgan of the IBERS Translational Genomics Facility.

---

**Figure S1.**
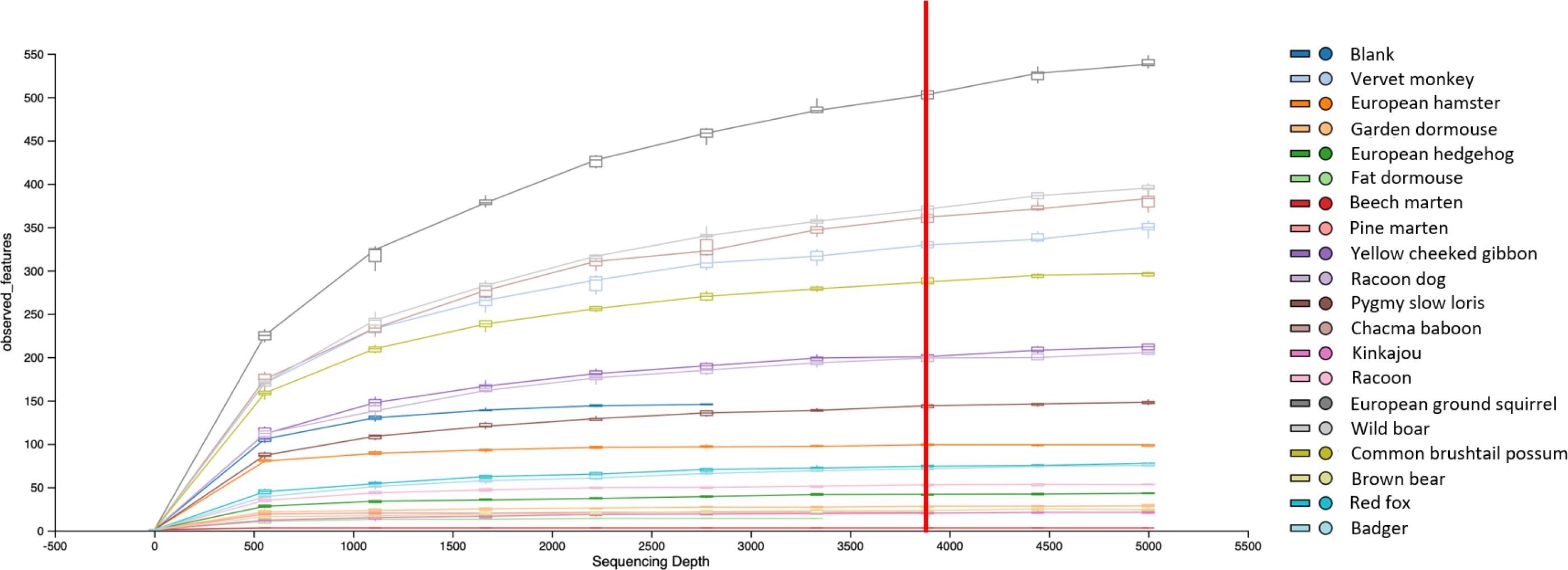
Rarefaction curve - Number of observed features at different sampling depths Sampling depth of 3816 marked with red line

**Figure S2.**
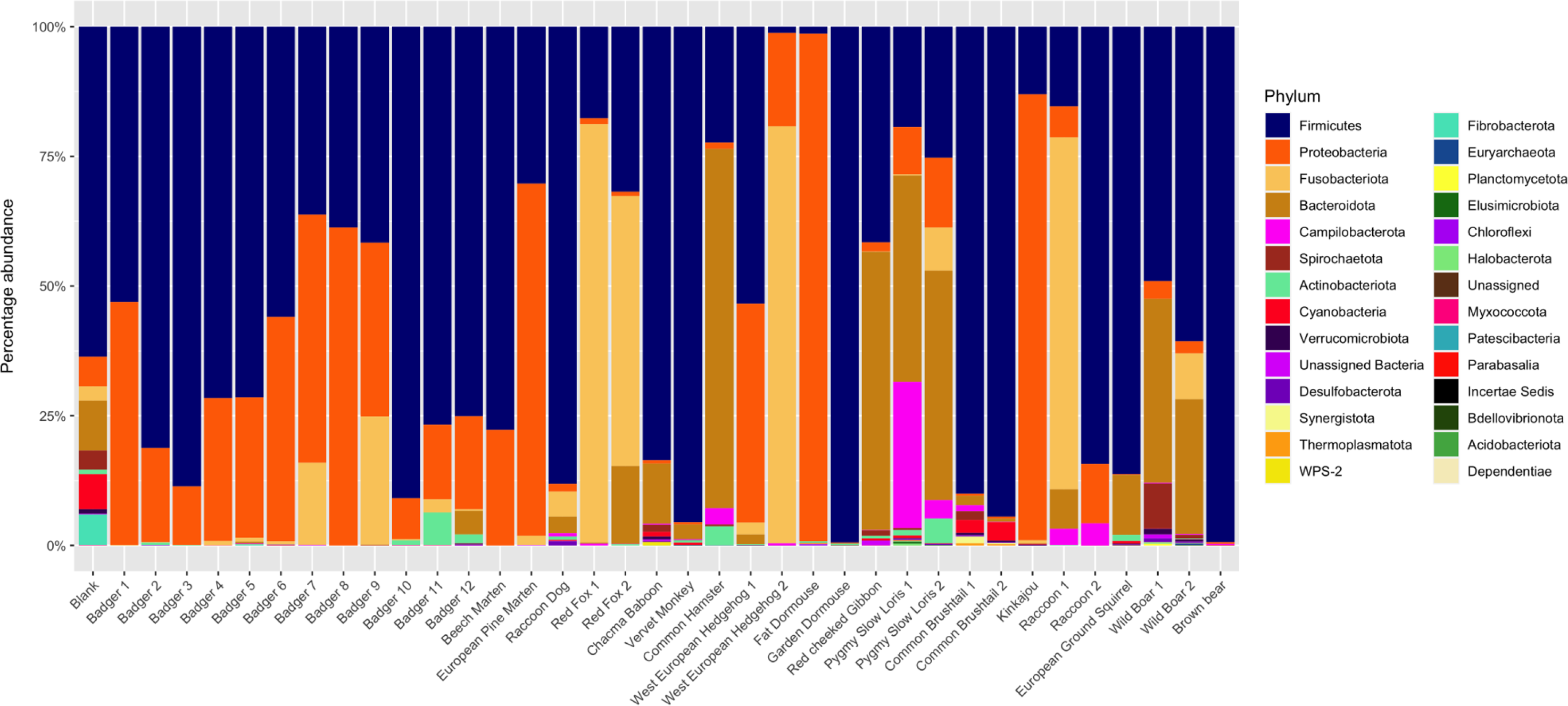
Assembling the omnivorous mammalian faecal microbiome collection: Percentage abundances of different phyla of bacteria

## Bibliography

Abernethy, D.A., Walton, E., Menzies, F., Courcier, E., and Robinson, P. (2011) *Mycobacterium bovis* surveillance in European badgers (*Meles meles*) killed by vehicles in Northern Ireland: an epidemiological evaluation. Int Conf Anim Heal Surveill 78: 216–218.

Bokulich, N.A., Kaehler, B.D., Rideout, J.R., Dillon, M., Bolyen, E., Knight, R., et al. (2018) Optimizing taxonomic classification of marker-gene amplicon sequences with QIIME 2’s q2-feature-classifier plugin. Microbiome 6: 90.

Bolyen, E., Rideout, J.R., Dillon, M.R., Bokulich, N.A., Abnet, C.C., Al-Ghalith, G.A., et al. (2019) Reproducible, interactive, scalable and extensible microbiome data science using QIIME 2. Nat Biotechnol 37: 852–857.

Brice, K., Trivedi, P., Jeffries, T., Blyton, M., Mitchell, C., Singh, B., and Moore, B. (2019) The Koala (*Phascolarctos cinereus*) faecal microbiome differs with diet in a wild population. PeerJ 7: e6534.

Choo, J.M., Leong, L.E.X., and Rogers, G.B. (2015) Sample storage conditions significantly influence faecal microbiome profiles. Sci Rep 5: 16350.

Clayton, J.B., Vangay, P., Huang, H., Ward, T., Hillmann, B.M., Al-Ghalith, G.A., et al. (2016) Captivity humanizes the primate microbiome. Proc Natl Acad Sci 113: 10376 LP – 10381.

Colston, T. and Jackson, C. (2016) Microbiome Evolution Along Divergent Branches of the Vertebrate Tree of Life: What’s Known and Unknown. Mol Ecol 25:.

Conlon, M. and Bird, A. (2014) The Impact of Diet and Lifestyle on Gut Microbiota and Human Health. Nutrients 7: 17–44.

Defra (2020) Next steps for the strategy for achieving bovine tuberculosis free status for England.

Deusch, O., O’Flynn, C., Colyer, A., Swanson, K.S., Allaway, D., and Morris, P. (2015) A Longitudinal Study of the Feline Faecal Microbiome Identifies Changes into Early Adulthood Irrespective of Sexual Development. PLoS One 10: e0144881.

Eckburg, P., Bik, E., Bernstein, C., Purdom, E., Dethlefsen, L., Sargent, M., et al. (2005) Diversity of the Human Intestinal Microbial Flora. Science 308: 1635–1638.

Enaud, R., Prevel, R., Ciarlo, E., Beaufils, F., Wieërs, G., Guery, B., and Delhaes, L. (2020) The Gut-Lung Axis in Health and Respiratory Diseases: A Place for Inter-Organ and Inter-Kingdom Crosstalks. Front Cell Infect Microbiol 10: 1–11.

Fava, F., Gitau, R., Griffin, B.A., Gibson, G.R., Tuohy, K.M., and Lovegrove, J.A. (2013) The type and quantity of dietary fat and carbohydrate alter faecal microbiome and short-chain fatty acid excretion in a metabolic syndrome ‘at-risk’ population. Int J Obes 37: 216–223.

Gibson, K.M., Nguyen, B.N., Neumann, L.M., Miller, M., Buss, P., Daniels, S., et al. (2019) Gut microbiome differences between wild and captive black rhinoceros – implications for rhino health. Sci Rep 9: 7570.

Gimmel, A., Eulenberger, U., and Liesegang, A. (2021) Feeding the European hedgehog (*Erinaceus europaeus* L.)—risks of commercial diets for wildlife. J Anim Physiol Anim Nutr (Berl) n/a:

Godfray, C., Donnelly, C., Hewinson, G., Winter, M., and Wood, J. (2018) Bovine TB Strategy Review.

Goodchild, A. V., Watkins, G.H., Sayers, A.R., Jones, J.R., and Clifton-Hadley, R.S. (2012) Geographical association between the genotype of bovine tuberculosis in found dead badgers and in cattle herds. Vet Rec 170: 259.

Grond, K., Sandercock, B., Jumpponen, A., and Zeglin, L. (2018) The Avian Gut Microbiota: Community, Physiology and Function in Wild Birds. J Avian Biol 49:.

Heinritz, S.N., Weiss, E., Eklund, M., Aumiller, T., Louis, S., Rings, A., et al. (2016) Intestinal Microbiota and Microbial Metabolites Are Changed in a Pig Model Fed a High-Fat/Low-Fiber or a Low-Fat/High-Fiber Diet. PLoS One 11: e0154329.

Hooda, S., Boler, B.M.V., Serao, M.C.R., Brulc, J.M., Staeger, M.A., Boileau, T.W., et al. (2012) 454 Pyrosequencing Reveals a Shift in Fecal Microbiota of Healthy Adult Men Consuming Polydextrose or Soluble Corn Fiber. J Nutr 142: 1259–1265.

Hooda, S., Vester Boler, B.M., Kerr, K.R., Dowd, S.E., and Swanson, K.S. (2013) The gut microbiome of kittens is affected by dietary protein:carbohydrate ratio and associated with blood metabolite and hormone concentrations. Br J Nutr 109: 1637–1646.

Hu, Y., Feng, Y., Wu, J., Liu, F., Zhang, Z., Hao, Y., et al. (2019) The gut microbiome signatures discriminate healthy from pulmonary tuberculosis patients. Front Cell Infect Microbiol 9: 1–8.

Hungerford, L.L., Mitchell, M.A., Nixon, C.M., Esker, T.E., Sullivan, J.B., Koerkenmeier, R., and Marretta, S.M. (1999) Periodontal and dental lesions in raccoons from a farming and a recreational area in Illinois. J Wildl Dis 35: 728–734.

Lees, H., Swann, J., Poucher, S.M., Nicholson, J.K., Holmes, E., Wilson, I.D., and Marchesi, J.R. (2014) Age and Microenvironment Outweigh Genetic Influence on the Zucker Rat Microbiome. PLoS One 9: e100916.

Ley, R., Hamady, M., Lozupone, C., Turnbaugh, P., Ramey, R., Bircher, J., et al. (2008) Evolution of Mammals and Their Gut Microbes. Science 320: 1647–1651.

Luo, M., Liu, Y., Wu, P., Luo, D.X., Sun, Q., Zheng, H., et al. (2017) Alternation of gut microbiota in patients with pulmonary tuberculosis. Front Physiol 8:.

Middelbos, I.S., Vester Boler, B.M., Qu, A., White, B.A., Swanson, K.S., and Fahey, G.C. Jr. (2010) Phylogenetic Characterization of Fecal Microbial Communities of Dogs Fed Diets with or without Supplemental Dietary Fiber Using 454 Pyrosequencing. PLoS One 5: 1–9.

Nelson, T., Apprill, A., Mann, J., Rogers, T., and Brown, M. (2015) The marine mammal microbiome: current knowledge and future directions. Microbiol Aust.

Ning, Y., Qi, J., Dobbins, M.T., Liang, X., Wang, J., Chen, S., et al. (2020) Comparative Analysis of Microbial Community Structure and Function in the Gut of Wild and Captive Amur Tiger. Front Microbiol 11: 1665.

Panasevich, M.R., Kerr, K.R., Dilger, R.N., Fahey, G.C., Guérin-Deremaux, L., Lynch, G.L., et al. (2015) Modulation of the faecal microbiome of healthy adult dogs by inclusion of potato fibre in the diet. Br J Nutr 113: 125–133.

Pechal, J.L., Schmidt, C.J., Jordan, H.R., and Benbow, M.E. (2018) A large-scale survey of the postmortem human microbiome, and its potential to provide insight into the living health condition. Sci Rep 8: 5724.

Potrykus, J., Povitz, B., White, R., and Bearne, S. (2007) Proteomic investigation of glucose metabolism in the butyrate-producing gut anaerobe *Fusobacterium varium*. Proteomics 7: 1839–1853.

Proudman, C., Hunter, J.O., Darby, A.C., Escalona, E.E., Batty, C., and Turner, C. (2014) Characterisation of the faecal metabolome and microbiome of Thoroughbred racehorses. 47: 580–586.

Quast, C., Pruesse, E., Yilmaz, P., Gerken, J., Schweer, T., Yarza, P., et al. (2013) The SILVA ribosomal RNA gene database project: improved data processing and web-based tools. Nucleic Acids Res 41: D590–D596.

Robertson, A., Mcdonald, R., Delahay, R., Kelly, S., and Bearhop, S. (2014) Individual foraging specialisation in a social mammal: the European badger (*Meles meles*). Oecologia 176:.

Robeson, M.S., O’Rourke, D.R., Kaehler, B.D., Ziemski, M., Dillon, M.R., Foster, J.T., and Bokulich, N.A. (2020) RESCRIPt: Reproducible sequence taxonomy reference database management for the masses. bioRxiv 2020.10.05.326504.

Sandoval Barron, E., Swift, B., Chantrey, J., Christley, R., Gardner, R., Jewell, C., et al. (2018) A study of tuberculosis in road traffic-killed badgers on the edge of the British bovine TB epidemic area. Sci Rep 8: 1–8.

Schmidt, E., Mykytczuk, N., and Schulte-Hostedde, A. (2019) Effects of the captive and wild environment on diversity of the gut microbiome of deer mice (P*eromyscus maniculatus*). ISME J 13:.

Schroeder, P., Hopkins, B., Jones, J., Galloway, T., Pike, R., Rolfe, S., and Hewinson, G. (2020) Temporal and spatial Mycobacterium bovis prevalence patterns as evidenced in the All Wales Badgers Found Dead (AWBFD) survey of infection 2014–2016. Sci Rep 10: 1–11.

Schulte-Hostedde, A.I., Mazal, Z., Jardine, C.M., and Gagnon, J. (2018) Enhanced access to anthropogenic food waste is related to hyperglycemia in raccoons (*Procyon lotor*). Conserv Physiol 6:.

Shanmuganandam, S., Hu, Y., Strive, T., Schwessinger, B., and Hall, R. (2020) Uncovering the microbiome of invasive sympatric European brown hares and European rabbits in Australia. PeerJ 8: e9564.

Sidorovich, V.E., Sidorovich, A.A., and Izotova, I. V (2006) Variations in the diet and population density of the red fox *Vulpes vulpes* in the mixed woodlands of northern Belarus. Mamm Biol 71: 74–89.

Thomas, R. and Chambers, M. (2021) Review of Methods Used for Diagnosing Tuberculosis in Captive and Free-Ranging Non-Bovid Species (2012–2020). Pathog 10:.

Wan, Y., Wang, F., Yuan, J., Li, J., Jiang, D., Zhang, J., et al. (2019) Effects of dietary fat on gut microbiota and faecal metabolites, and their relationship with cardiometabolic risk factors: a 6-month randomised controlled-feeding trial. Gut 68: 1417–1429.

Wu, G., Chen, J., Hoffmann, C., Bittinger, K., Chen, Y.-Y., Keilbaugh, S., et al. (2011) Linking Long-Term Dietary Patterns with Gut Microbial Enterotypes. Science 334: 105–108.

Youngblut, N.D., Reischer, G.H., Walters, W., Schuster, N., Walzer, C., Stalder, G., et al. (2019) Host diet and evolutionary history explain different aspects of gut microbiome diversity among vertebrate clades. Nat Commun 10: 2200.

